# Molecular mechanism of α-synuclein aggregation on lipid membranes revealed

**DOI:** 10.1101/2023.10.20.563279

**Authors:** Alexander J. Dear, Xiangyu Teng, Sarah R. Ball, Joshua Lewin, Robert I. Horne, Daniel Clow, Natasha Harper, Kim Yahya, Thomas C.T. Michaels, Sara Linse, Tuomas P. J. Knowles, Xiaoting Yang, Suzanne C. Brewerton, John Thomson, Johnny Habchi, Georg Meisl

## Abstract

The central hallmark of Parkinson’s disease pathology is the aggregation of the *α*-synuclein protein, which, in its healthy form, is associated with lipid membranes. Purified monomeric *α*-synuclein is relatively stable in vitro, but its aggregation can be triggered by the presence of lipid vesicles. Despite this central importance of lipids in the context of *α*-synuclein aggregation, their mechanistic role in this process has not been established to date. Here, we use chemical kinetics to develop a detailed mechanistic model that is able to globally describe the aggregation behaviour of *α*-synuclein in the presence of DMPS lipid vesicles, across a range of lipid and protein concentrations. Through the application of our kinetic model to experimental data, we find that the reaction is a co-aggregation process involving both protein and lipids and that lipids promote aggregation predominantly by enabling the elongation process. Moreover, we find that the initial formation of aggregates, via primary nucleation, takes place not on the surface of lipid vesicles but at the interfaces present in vitro. Our model will enable mechanistic insights, also in other lipid-protein co-aggregation systems, which will be crucial in the rational design of drugs that inhibit aggregate formation and act at the key points in the *α*-synuclein aggregation cascade.

## INTRODUCTION

The aggregation of *α*-synuclein has been linked to the emergence of a range of neurodegenerative disorders[1– 3], the synucleinopathies, the most prominent of which is Parkinson’s disease. Thus, *α*-synuclein aggregation is a promising target for drug development and significant efforts have been made to discover inhibitors of this process[4–7]. A key requirement for the successful development of small molecule aggregation inhibitors is the availability of a reliable, predictive in vitro assay, which often relies on a purified protein drug target, to evaluate compound potency. While this parallels drug discovery strategies for other targets, the search for aggregation inhibitors is additionally complicated by the complexity of the aggregation reaction: several different steps contribute to the overall aggregate formation reaction[8, 9], and all are potential targets for slowing in vitro aggregation. However, targeting some of these different steps is unlikely to elicit the desired in vivo responses[10–12]. A detailed mechanistic understanding of the aggregation mechanism and its inhibition is thus required for the accurate interpretation of in vitro data[13].

In recent decades, mechanistic work has shown that the aggregation of most purified proteins into large fibrillar aggregates in vitro involves at least 3 classes of processes [15, 16]: *primary nucleation*, which leads to the formation of new aggregates directly from monomeric protein without the involvement of existing aggregates, *secondary processes* (such as fragmentation or secondary nucleation), which lead to the formation of new aggregates from existing aggregates and *elongation*, which is the growth of existing aggregates by addition of monomeric protein. By applying the framework of chemical kinetics[8], these mechanisms can be turned into rate laws to then be fitted to experimental data[13]. Application of these rate laws to measurements of the aggregation of purified protein in vitro has been very successful in elucidating the detailed mechanisms of aggregate formation for a wide range of proteins[16], in particular A*β*[17–19], but to some extent also *α*-synuclein under different conditions[20, 21]. However, the more complex mechanism that describes the aggregation of a mixture of purified protein and lipid vesicles has remained elusive to date. Here, we build such a mechanistic description and show that it can describe the aggregation of *α*-synuclein and 1,2-dimyristoyl-*sn*-glycero-3-phospho-L-serine (DMPS) lipid vesicles, globally, across lipid and protein concentrations.

At neutral pH, purified *α*-synuclein is surprisingly resistant to aggregation, generally requiring vigorous agitation or the introduction of preformed seed fibrils to trigger the aggregation process[20, 22]. This resistance of *α*-synuclein to aggregation is likely due to the low rate of primary nucleation. The barrier to nucleation is high, meaning monomeric *α*-synuclein is kinetically stable, but agitation, likely by introducing shearing forces and increasing turnover at the air-water interface can significantly increase its rate[23]. Introduction of preformed aggregates increases the speed of aggregation considerably, further supporting the idea that a slow primary nucleation step is the main reason for the aggregation resistance of *α*-synuclein[20]. Lowering the pH [20, 21] or using high salt concentrations [24] are other ways to induce aggregation, likely by reducing electrostatic repulsion between the aggregating proteins[25, 26].

An alternative method to initiate the aggregation at neutral pH is the introduction of lipids[27], usually in the form of small unilamellar vesicles (SUVs)[28], see Fig. 1, which induces formation of aggregates without extended lagtimes. Here we focus on lipid vesicles made from DMPS lipids, but similar effects have been observed with a range of lipid compositions, suggesting a general behaviour[27, 29]. DMPS lipids have short saturated acyl chains and its membranes have a melting temperature above 37^*o*^C [30]. Although not found in biological membranes, DMPS in the form of SUVs efficiently triggers α-synuclein aggregation at neutral pH and may therefore serve as a useful system for mechanistic in vitro studies, such as inhibitor screening.

**FIG. 1.**
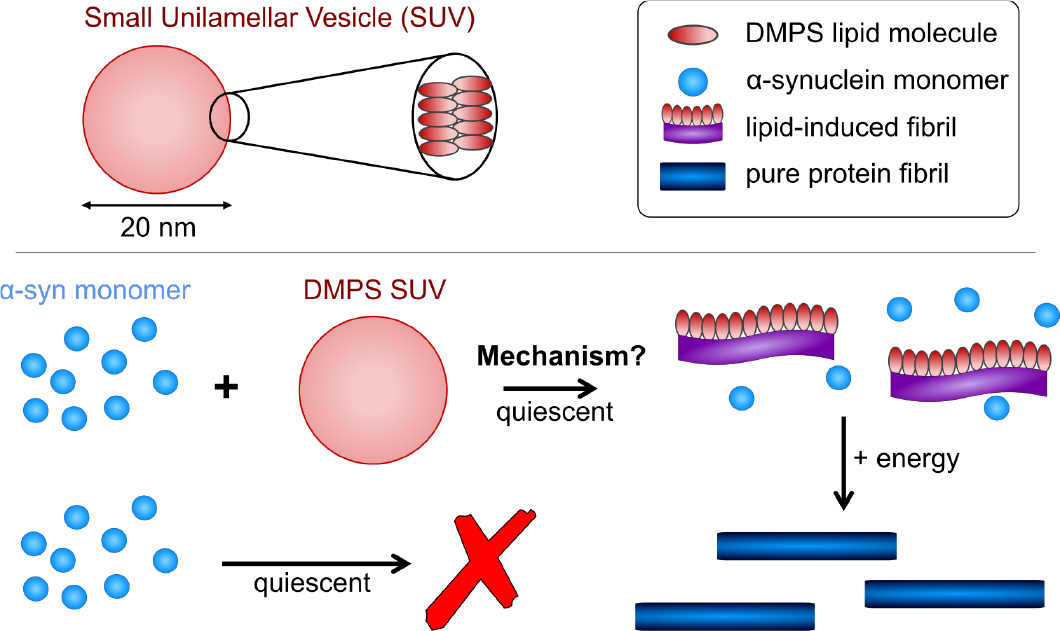
Schematic of *α*-synuclein aggregates w/o lipids. In the presence of lipid vesicles *α*-synuclein forms mixed protein-lipid aggregates. In the absence of lipids, monomeric *α*-synuclein is much more stable and biased against aggregation, however, the formation of pure protein fibrils can be triggered e.g. by agitation. Lipidic fibrils can similarly be triggered to convert to the structures seen in pure protein aggregation e.g. by heating the reaction mixture[14].

In vivo, *α*-synuclein is believed to be associated with lipid membranes, potentially as part of both its biological function and its pathology[31–35]. The introduction of lipid vesicles thus not only allows formation of aggregates under quiescent conditions in vitro, but it also provides a simple in vitro model for the interaction of *α*-synuclein with membranes. The interaction of *α*-synuclein with lipid-vesicles and their effect on the kinetics of aggregation has been investigated in detail [14, 27, 28, 32, 34–38]. It was thus established, using model membranes, that *α*-synuclein adsorbs in an *α*-helical conformation [39] in the head group area and upper acyl layer [40, 41] and that protein aggregation is triggered only in situations of protein excess [28, 29, 42]. However, to date no mechanistic model has been able to globally describe the observed aggregation kinetics and thus establish how and at which microscopic step lipid vesicles promote the aggregation of *α*-synuclein.

While DMPS lipid vesicles efficiently trigger the aggregation of *α*-synuclein, the fibrils formed under those conditions differ considerably from those formed by seeding or through agitation[14]. Cryo-EM images reveal that the aggregates formed in the presence of DMPS [30] are also different from those formed in the presence of other lipids such as DOPC-DOPS mixtures [43] and DOPC-GM1 mixtures [44], although lipids appear to decorate the fibrils in all cases. The fibrils formed in the presence of DMPS lipids are sometimes referred to as protofibrils with those formed without lipids referred to as mature fibrils. To minimise the potential for misinterpretation, we here refer to them as lipidic fibrils and pure protein fibrils, respectively. While pure protein fibrils form long, straight structures, the lipidic fibrils appear more flexible in microscopy[14, 28]. There is strong evidence that lipidic fibrils are in fact co-aggregates of both *α*-synuclein and lipids[28, 30], not just for DMPS but across different lipid systems [43], with some recent work providing high resolution structures of such mixed fibrils[45]. Another strong piece of evidence in favour of lipidic fibrils containing both lipid and protein comes from the fact that the amount of lipidic fibrils formed is limited not only by the amount of available protein, but also by the amount of available lipid, see Fig. 2A. It has been shown that the structures for lipidic and pure protein fibrils are, however, not entirely unrelated: heating of lipidic fibrils can induce conversion to pure protein fibrils, indicating that the lipidic fibrils are a thermodynamically less stable form[14]. This observation has also given rise to the hypothesis that some protein-lipid co-aggregates represent the precursors to mature aggregates in disease, prompting our research into therapeutic molecules to target this process. Therefore, a mechanistic understanding of exactly how lipids induce *α*-synuclein aggregation is crucial both to evaluate the translatability of in vitro lipid-induced experiments to in vivo systems and to establish which processes in this aggregation reaction are rate-limiting and therefore should be the focus of drug development efforts. In this work, we focus on DMPS lipids and present a new kinetic model that is able to globally fit the aggregation kinetics, across monomer and DMPS lipid concentrations, and thus yields new insights into the ways in which lipid vesicles do (or do not) trigger *α*-synuclein aggregation.

**FIG. 2.**
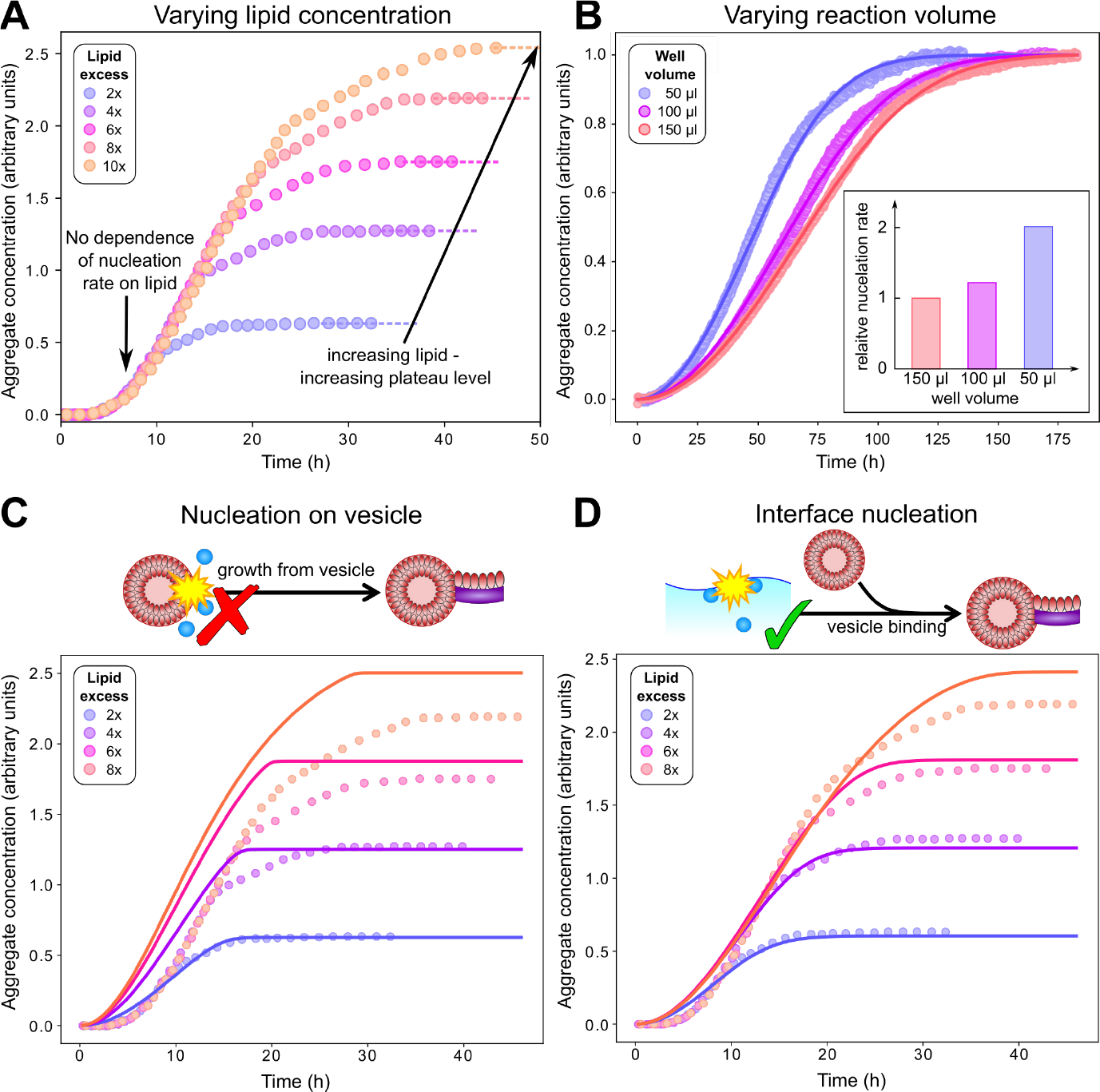
Kinetic analysis shows that primary nucleation does not occur on vesicles. **A**: Careful inspection of kinetic data from Galvagnion et al. [28] shows that aggregation rates of *α*-synuclein (50 _*μ*_M) are independent of lipid concentration before the plateauing of aggregation curves. **B**: Aggregation experiments (20 _*μ*_M *α*-synuclein + 40 _*μ*_M DMPS) with varying surface area to volume ratio show that primary nucleation rates increase with this ratio (fits to Eq. (1)), confirming that primary nucleation is heterogeneous. **C**: A kinetic model featuring primary nucleation on vesicles cannot fit the data from A. **D**: Conversely, a model featuring primary nucleation on reaction vessel interfaces Eq. (1)) can fit the data reasonably well (the plateau heights can only be captured approximately for reasons discussed later; see later sections for model details; data in A,C,D from Galvagnion et al.[28]). Parameters: 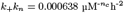, *n*_*c*_ = 0.6, *k*_2_ = 0, *y* = 470, *k*_on_*/k*_+_ = 9.5.

## RESULTS AND DISCUSSION

The prevailing perception is that primary nucleation takes place on the surface of vesicles, for example through the interaction of free protein with protein bound to vesicles, based on the fact that without lipids, the lack of primary nucleation prevents aggregate formation whereas in the presence of lipids aggregate formation is observed[28]. However, while compelling at first, closer examination of the aggregation of α-synuclein in the presence of DMPS lipids shows that this conclusion does not necessarily follow and there are a number of equally simple mechanistic explanations for the observed kinetics. First, we clarify the terminology used here: in the presence of DMPS lipids, the primary aggregated species are lipidic, rather than pure protein fibrils. It is the processes that give rise to these lipidic fibrils, i.e. their nucleation, growth and potentially fragmentation / secondary nucleation which we here investigate. If one were to consider a system in which the conversion of lipidic fibrils to pure protein fibrils is induced, for example by a temperature jump[14], lipidic fibrils can be the precursors to pure protein fibrils. In such a system, lipid-induced aggregation can be considered as an initiation event for the formation of pure protein fibrils. However, we feel that considering the entirety of the formation, growth and conversion of lipidic fibrils as the “primary nucleation” step of pure protein fibril formation is stretching the definition of nucleation beyond its generally accepted meaning, so will not be using this terminology. Thus, the question becomes, in which steps of the formation of lipidic fibrils does the presence of lipids play a key role? We will now examine these processes in turn and build a kinetic model to globally fit the aggregation kinetics across lipid and monomer concentrations.

It is worth noting that, because *α*-synuclein monomers bind to lipid vesicles, at high lipid to protein ratios essentially all protein will be bound to vesicles, very little protein remains in solution and no aggregation takes place[28]. In this work, we will only consider situations in which a significant excess of protein is present, so lipid vesicles are saturated in bound protein and the majority of protein is present in solution, not in its lipid bound form, allowing for aggregate formation.

### Primary nucleation does not take place on the surface of lipid vesicles

First we investigate the origin of the initial fibrils via primary nucleation. If indeed primary nucleation were a simple heterogeneous nucleation process on the surface of lipid vesicles, with or without the involvement of both lipid bound and free protein, the data would be expected to display a number of key features. Most importantly one would expect the rate of primary nucleation to depend on the concentration of lipid vesicles present. In this model the vesicle surface serves as a catalytic site for monomer conversion, so the dependence of the rate on vesicle concentration would be linear as an increase in the vesicle concentration simply corresponds to an increase in catalytic surface area. However, in reality, no such dependence of the rate on lipid concentration is observed, as demonstrated in Fig. 2**A**, where the aggregation reaction at a constant α-synuclein concentration and varying DMPS lipid concentrations is monitored. In this experiment, the kinetic curves at different lipid concentrations overlap perfectly until the plateau phase begins. This observation suggests not only that nucleation does not occur on DMPS vesicles, but that before the plateau phase, lipid involvement is not rate-limiting in any of the aggregation processes, at the lipid and monomer concentrations used. The lipid concentration only becomes limiting as the reaction nears its plateau, with higher lipid concentrations leading to higher plateau levels.

If primary nucleation does not occur on vesicles, the remaining possibilities are that it is a homogeneous process occurring in solution, or that it occurs on the air-water interface or the plate surface. To further experimentally evaluate the importance of interface effects, we monitored aggregation reactions with the same starting monomer and lipid concentrations, but with different solution volumes of 50, 100 and 150 *μ*l, to alter the surface to volume ratio (Fig. 2B). We observed a significant increase in the rate of aggregation as the volume decreased (and thus the surface area to volume ratio increased), consistent with primary nucleation being a heterogeneous, surface catalysed process. This is in line with the general observation that heterogeneous, rather than homogeneous, primary nucleation is likely the more common process in most aggregating systems [23, 46]. This finding is confirmed by the misfit or fit of a model assuming nucleation on vesicles or on interfaces, respectively, see Fig. 2C and D (for details on the other steps in the model see below).

While these data indicate the involvement of interfaces in nucleation and rule out primary nucleation taking place directly on the surface of DMPS vesicles, they do not allow us to determine whether lipids are involved in the nucleation step at the interfaces. Lipid involvement would be consistent with both the lack of a lipid concentration dependence and the sensitivity to the surface to volume ratio, if lipids fully cover the interface on which heterogeneous nucleation takes place. Similarly, a nucleation reaction not involving any lipid would display the same kinetic signature, thus preventing clear distinction of the two mechanisms based on the present kinetic data. However, considering the fact that lipids typically form a monolayer at the air water interface, with the head group facing the water and the acyl chains the air, and the observation that lipid-protein co-aggregates are formed, lipid involvement also during nucleation seems likely.

### Presence of lipid is crucial for elongation of lipidic fibrils

Having demonstrated that primary nucleation predominantly does not take place on DMPS vesicles, and is its rate is independent of lipid concentration, the question remains how DMPS lipids induce aggregate formation. In this context, it is crucial to appreciate how essential fibril elongation is for the observation of any aggregate formation in an amyloid forming system: while primary nucleation, and in some systems secondary nucleation, are responsible for the formation of new fibrils, these new fibrils are usually orders of magnitude smaller than the average fibril size observed at the end of the reaction. Fibrils generally contain thousands to tens of thousands of protein monomers, whereas newly nucleated fibrils likely contain only tens of monomers. Thus, for every nucleation event thousands of elongation events take place[13, 15, 16]. While introduction of preformed seeds enables aggregation of a monomeric sample by bypassing nucleation, elongation cannot be bypassed and without elongation, there will be no aggregation.

The key observation is that fibrils formed in the presence of DMPS vesicles are co-aggregates of protein and lipids and that the amount of fibrillar material appears to be limited by the amount of lipids present (when protein is present in excess). Thus, even at the plateau of the aggregation curve, significant amounts of protein monomer remain in solution. Taken together these findings indicate that DMPS lipids are either directly involved in the elongation step or that they have to stabilize newly extended fibrils soon after monomer addition. In Fig. 3**A** we show that the relative fibril mass concentration formed by the end of the reaction scales linearly with lipid concentration, across several different experiments, at a range of protein concentrations. Note that at even higher lipid concentrations this linear relationship will break down, once the availability of protein, not lipid, limits the amount of fibrils formed. With these insights, we can now build a mechanistic model to globally fit the aggregation kinetics[13, 15].

**FIG. 3.**
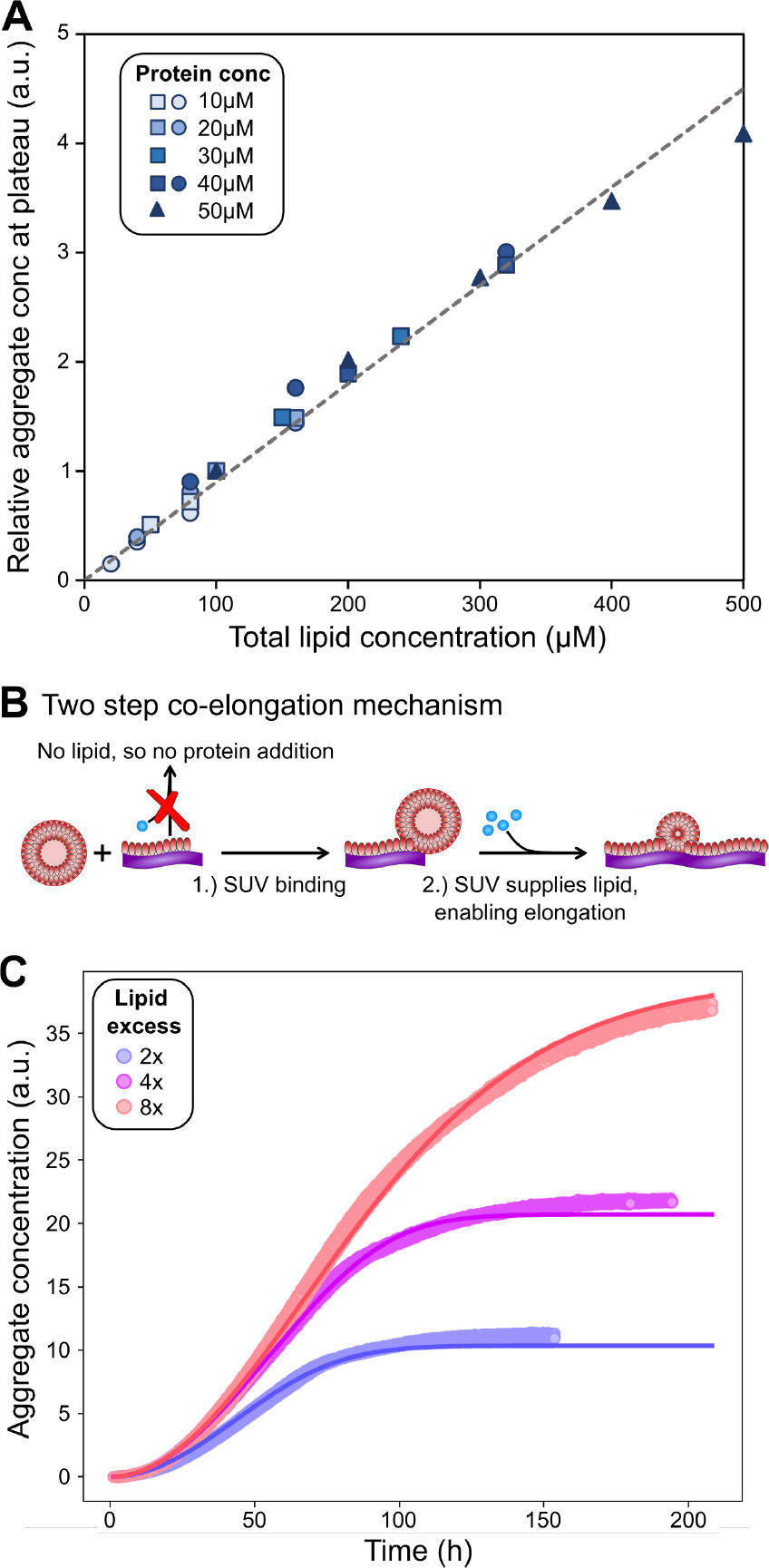
Lipids are involved in co-elongation of fibrils. **A** Amounts of fibril at plateau of aggregation curves, as reported by ThT intensity, scale linearly with lipid amounts. Several datasets, at a range of different monomer concentrations, were combined to generate this plot. Circles and squares are new data, triangles are the data from Galvagnion et al.[28]. To account for differences in ThT fluorescence between the three datasets, within each set the data were normalised to the ThT intensity at 100*μ*M (80 *μ*M for circles) lipid. **B** Schematic of lipid-protein co-elongation reaction. **C** Aggregation at a constant monomer (40 *μ*M) and varying lipid concentrations (2x, 4x and 8x), fitted to a model of surface catalysed primary nucleation and lipid-protein co-elongation, see Equ. 1.

### Building a kinetic model of lipid-induced aggregation

Before a kinetic description of fibril formation is possible, it is necessary to determine which species are formed over the course of the reaction. Initially, only DMPS vesicles and monomeric *α*-synuclein are present. However, monomeric *α*-synuclein binds tightly to the vesicles, a process that is many times faster than the tens of hours it takes for lipidic fibrils to form[28, 31, 38]. For kinetic modelling of the fibril formation process, it is therefore acceptable to approximate the *α*-synuclein monomer-vesicle binding as pre-equilibrium, i.e. assume this equilibrium has already been attained by *t* = 0. Using the previously determined stoichiometry and affinity of *α*-synuclein to lipid vesicles[28], we can compute the remaining free *α*-synuclein concentration. Under the conditions used in this work, *α*-synuclein is always present in significant excess, such that vesicles are fully covered and the majority of *α*-synuclein is in solution at the beginning of the reaction.

At the end of the aggregation reaction, there is an equilibrium of lipidic fibrils, monomeric *α*-synuclein, free vesicles and vesicle-bound *α*-synuclein. However, we can simplify the picture by setting the concentration of free vesicles to zero, due to the excess protein and the tight binding. The remaining unknown is the stoichiometry of *α*-synuclein and DMPS in lipidic fibrils. It can be estimated from measurements of fibril formation, under a varying initial lipid:protein ratio *r*_0_ and at a fixed initial *α*-synuclein concentration. As shown previously[28], the yield of lipidic fibrils, as reported by the height of the plateau of the kinetic curves, reaches a maximum at *r*_0_ *≃* 15, implying that the optimal stoichiometry of lipid:protein in lipidic fibrils is approximately 15. This can be rationalized as follows. If the optimal lipid:protein stoichiometry in the fibrils is *x*, then when *r*_0_ *< x*, as fibril formation progresses the ratio of lipid:protein outside the fibrils decreases as the lipid is used up faster in relative terms. Thus the yield is ultimately lipid-limited. The inverse argument applies when *r*_0_ *> x*, where protein is depleted faster and the yield is protein-limited. Thus, the ratio of initial protein and lipid concentrations at the maximum yield gives the the ratio in the lipidic fibrils as *x* = *r*_0_ *≃* 15. This estimate for the stoichiometry in lipidic fibrils is used in the remainder of this work.

To build the kinetic model, we first specify the rates of all processes involved, adhering to previous convention where possible[8, 13, 15]. The rate of primary nucleation is given by 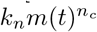, where *m*(*t*) is the *α*-synuclein monomer concentration. A value of *n*_*c*_ below 1 indicates saturation of the interface where primary nucleation is occurring[46]. The concentration of heterogeneous nucleation sites is subsumed into the rate constant *k*_*n*_. As no dependence of the primary nucleation rate on the DMPS lipid concentration is observed, this is not modelled explicitly. A potential secondary nucleation rate can be written as 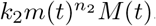*M* (*t*), where *M* (*t*) is the *α*-synuclein fibril concentration, *k*_2_ is the rate constant and *n*_2_ the reaction order with respect to protein.

Elongation, involving addition of both protein and lipid to form a lipid-protein co-aggregate, is more complicated to model. We approach it as follows: First, a lipid vesicle (concentration *c*_*S*_) binds to the surface of a fibril adjacent to the growing end. This then enables the sequential addition of *y* monomers on average to the growing end, where *y* is limited by the extent to which the vesicle can provide lipids to the growing fibril, and can be related to the lipid to protein stoichiometry, *x*. Once the *y*-th monomer has been added, a new vesicle must bind adjacent to the newly positioned growing end before any further elongation by monomer addition may occur, see Fig. 3B.

These considerations lead to the following overall kinetic model (derived in Methods):

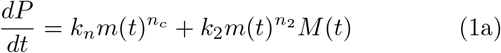

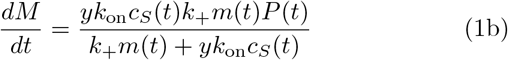

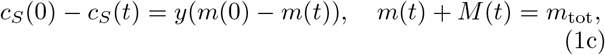

where *P* (*t*) is the total concentration of fibril ends, *k* _on_ is the rate constant for binding of vesicles to fibrils, and *k*_+_ the rate constant for elongation by addition of a protein monomer to the fibril end. A lthough t his non-covalent co-assembly is reversible, the back reactions are assumed to be significantly slower than the forward reactions and are therefore usually neglected in a kinetic model of fibril formation [15]. Fits of this model are shown in Fig. 3C.

A type of process that is believed to be key in many pathological aggregating system is fibril self-replication[16]. That is any processes by which new fibrils are formed from existing fibrils, such as fragmentation or secondary nucleation. Their presence gives rise to self-replication of aggregated structures and characteristic kinetic curves with a pronounced lag phase[16]. While such a process can be included in our model (see Fig 5), the improvement in fit quality was minor and judged to be not significant enough given the increased complexity of the model. This tells us that in this lipid-induced assay, secondary nucleation or fragmentation, if present at all, only has a relatively minor effect on the overall aggregation kinetics. This finding is in line with previous work showing that a different s et of conditions w as generally required to enable significant secondary nucleation[20, 22, 24].

**FIG. 4.**
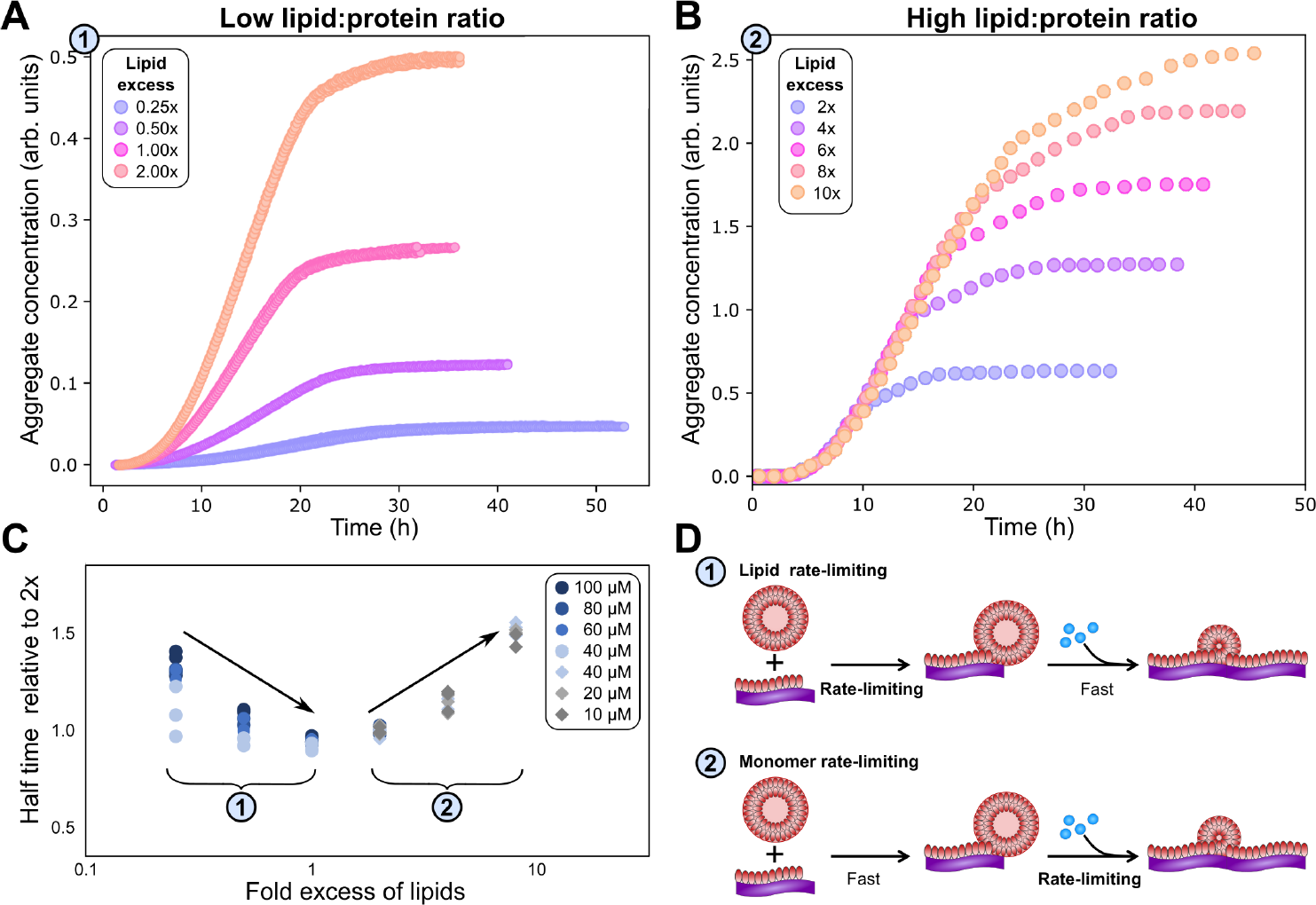
Lipid concentration limits rate at low lipid:protein ratios. **A** At lipid:protein ratios below 2, both the plateau height and the rate of aggregate formation depend on the lipid concentration. (*α*-synuclein monomer concentration is 100 _*μ*_M here.) **B** By contrast, at higher lipid:protein ratios the rate is independent of the lipid concentration (data in B is same as in Fig. 2A for direct comparison). **C** The half time of aggregation (time to reach half of the plateau aggregate concentration) is plotted against lipid:protein ratio. A number of different protein concentrations are collapsed onto the same curve by normalising the half times to that of the lipid:protein ratio of 2. Two regions in the data are clear from the half time plots, in which either the lipid or the protein is rate limiting. **D** Schematic mechanisms in the two regimes.

**FIG. 5.**
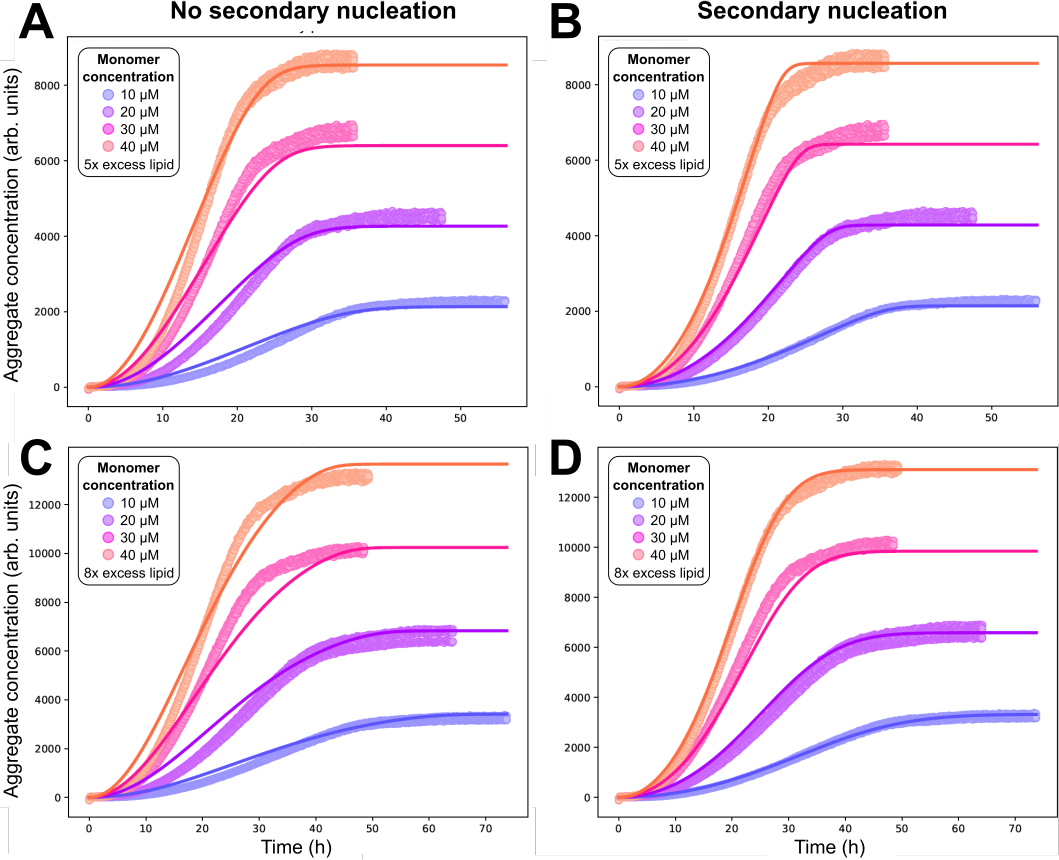
Kinetic model fitting to experimental data provides tentative evidence for weak secondary nucleation. **A**,**B**: 5x molar ratio of monomeric lipid to *α*-synuclein. **C**,**D**: 8x molar ratio of monomeric lipid to *α*-synuclein. **A**,**C**: setting *k*_2_ = 0 yields acceptable global fits of Eqs. (1) to the data. **B**,**D**: freely fitting *k*_2_ yields improved fits, but only slightly, implying a relatively slow secondary nucleation process may be present.

### Behaviour in lipid concentration limited regime confirms model

From the structure of Eq. (1b) it can be seen that when the rate of vesicle binding, *k*_on_, is large, its contribution becomes negligible. In that limit, the elongation rate depends only on protein monomer addition, reflecting that this has become the rate-limiting step for fibril elongation. Conversely, when the rate of protein monomer addition, *k*_+_, is large, it in turn becomes negligible, and vesicle binding becomes the rate-limiting step for fibril elongation.

The prediction that at low enough lipid concentrations vesicle attachment to the growing fibril should become rate-limiting can be tested experimentally[47]. Indeed, we find that below a DMPS lipid:protein ratio of 2, the rate of aggregation is reduced significantly when lipid concentration is reduced, see Fig. 4A, while at higher DMPS ratios the rate is constant, see Fig. 4B. This observation holds across different datasets and protein concentrations, as shown in Fig. 4C, where the half time of aggregation (the time at which half of the plateau aggregate mass is formed) is plotted against the lipid:protein ratio. The data acquired at different protein concentrations are collapsed onto one plot by rescaling the half times to the half time at a lipid:protein ratio of 2 (this is the condition shared across datasets). At ratios below 2, the half time decreases with increasing lipid concentration, i.e. the reaction speeds up with increasing lipid, which we refer to as regime 1. At ratios above 2, in regime 2, the half time increases with increasing lipid. Note that the reaction is not slower in absolute terms. This increase in half time simply results from the fact that the plateau increases with increasing lipid but the reaction proceeds at the same absolute speed, meaning it takes longer to reach the half way point between baseline and plateau.

These observations confirm that lipid is involved at the elongation step, rather than being added after fibrils have already formed, as concluded earlier based on plateau heights. However, they also allow us to rule out another possible mechanism, that of lipids attaching to growing fibrils as lipid monomers from solution, rather than as vesicles. While solubility of lipid monomers is low, some will be present in solution, in equilibrium with the more stable vesicle forms. Paralleling the behaviour in micelle formation, the lipid monomer concentration in solution is expected to be constant at lipid concentrations where vesicles are stable. As DMPS vesicles are stable at all lipid concentrations investigated here, we expect a constant concentration of DMPS monomers in solution. The fact that we observe the reaction rate to be dependent on lipid concentration is therefore in disagreement with a model where lipid addition proceeds by addition of lipid monomers from solution.

Finally, the fact that the same behaviour is observed for different protein concentrations when plotted against the lipid:protein ratio, confirms that the relative rates of protein and vesicle addition determine which regime the reaction is in.

### Implications for the design of inhibitors of aggregation

Having established the detailed mechanism of formation of lipidic fibrils, the key question is how this process can be inhibited most effectively, and which mode of action is most promising in the context of developing a drug against Parkinson’s disease[10, 13, 48]. Compounds targetting the initial formation of aggregates can be effective in vitro, for example by blocking the interfaces on which nucleation takes place, but are unlikely to translate to the in vivo situation where such interfaces may not be present in the same form[12]. Much more promising are compounds that inhibit aggregation by interacting with those processes expected to be relevant in vivo, in this case the incorporation of lipids or proteins into a growing fibril.

More generally, targetting fibrils is the more promising strategy for designing aggregation inhibitors for a number of reasons. Binding to fibril surfaces can inhibit fibril proliferation by preventing the formation of new fibrils via secondary nucleation, but it can also stop the production of toxic oligomeric species, directly and swiftly[11, 49]. Furthermore, it may constitute the safer approach in terms of avoiding inhibitor mechanism-based (ie. “on target”) toxicity. The aggregated structures are products of a disease process and appear to be greatly enriched in disease[2, 50]. Thus, binding to them is less likely to have deleterious effects when compared with binding to other targets that are known to be present and potentially important in healthy individuals, such as lipids or monomeric protein. This strategy is analogous to directly and specifically targeting the invading pathogen mechanisms in an infectious disease.

Our models show that, in the context of DMPS vesicle-induced aggregation of α-synuclein, targetting the incorporation of either monomer or lipid into the growing fibril are promising strategies for inhibiting aggregation in vitro. The above considerations highlight the fibril as the most suitable target for small molecules to inhibit this process and maximise chances of translatability. By contrast, given the negligible effect on the kinetics in this particular *in vitro* assay of any secondary processes that may be present, their inhibition will have little effect on the aggregation speed, and instead different conditions should be used to investigate this process[20, 22, 24].

## CONCLUSIONS

At neutral pH, in the absence of lipids, *α*-synuclein is kinetically stable, biased against aggregation for many hours due to the slow formation of pure protein aggregates. The observation that introduction of lipid vesicles triggers formation of aggregates has lead to the assumption that vesicles promote primary nucleation. However, our findings in this work reveal that the key effect of lipids in the case of DMPS vesicles is the promotion of fibril growth, likely by incorporation of lipids stabilising otherwise unstable protein aggregates. Quantitatively accounting for this formation of lipid-protein co-aggregates is sufficient to describe the observed kinetics, including the dependence of the aggregation rate on both protein and lipid concentrations. Primary nucleation of these lipidic fibrils may itself involve lipids, although the data are consistent also with primary nuclei being formed without lipid and lipids being added in a subsequent step. What can be ruled out is that the nucleation takes place on the surface of DMPS lipid vesicles, and we instead find that it is more likely that, in this in vitro system, primary nucleation occurs on the air-water or plate interfaces. The finding that primary nucleation takes place on these assay-dependent surfaces is an important point to consider when translating these finding between assays and drawing conclusions for the behaviour in living systems.

While the findings in this work are based on the behaviour of a specific lipid, DMPS, we expect that several aspects of it will translate to other lipids. In particular, if protein-lipid co-aggregates are formed as the predominant species, as seen with other lipids[43, 44], many of the considerations from this work will apply and the mechanism of elongation we propose here may also translate. Moreover, the air water interface or plate interfaces are found to be sites for primary nucleation across different proteins and experimental setups[23, 51], so our finding of interface-catalysed nucleation may be equally general. Finally, secondary nucleation appears to be only dominant under specific conditions in the aggregation of pure *α*-synuclein[20, 22, 24], so it would not be surprising if it is found to also be insignificant with lipids other than DMPS.

Beyond improving our mechanistic understanding, our findings highlight several key implications for the use of lipid-induced aggregation in the investigation of synucleinopathies and potential therapeutics: we show how lipidic fibrils differ from pure protein fibrils not only in their structure but also in their mechanism of formation. As both types of structure may be of relevance in disease, care needs to be taken when investigating mechanistic effects in vitro: potential therapeutic molecules should be capable of delaying fibril formation under both pure protein and lipid-induced conditions, to maximise their efficacy potential. The most robust way to achieve such dual pharmacology is to target the species that is key in both processes, the fibril surface. Moreover, we find that the DMPS vesicles provide lipid for the growth of lipidic fibrils, but they do not serve as nucleation sites, which instead takes place at air-water or plate interfaces. Thus, care needs to be taken when interpreting the readouts of this assay to ensure that the focus is on processes that retain in vivo translatability, and not on processes that involve air-water or plate interfaces.

In conclusion, the model we present here enables the quantitative analysis of DMPS *α*-synuclein co-aggregation, across protein and lipid concentrations. It is the first model of this kind and can serve as the basis for developing similar models in other systems where lipid-protein co-aggregation occurs. This mechanistic understanding of *α*-synuclein lipid co-aggregation will allow more targetted design of inhibitors of aggregation, that efficiently prevent the formation of both lipidic and pure protein fibrils and thus attack the formation of pathological aggregates at multiple points in the aggregation cascade.

## METHODS AND SUPPLEMENTARY INFORMATION

### Aggregation assay

The conditions in our lipid-induced aggregation assay are the same as those detailed in Galvagnion et al. [28], lipid and monomer concentrations vary and are specified at the relevant places in the text. The formation of pure protein fibrils is measured under the conditions detailed for the secondary nucleation assay in Buell et al. [20].

### Derivation of rate equations

Let *c*_*S*_(*t*) be the concentration of protein monomer-coated SUVs, and let *y* be the number of protein monomers (concentration *m*(*t*)) that add to the growing fibril by elongation on average each time an SUV binds to the fibril. Then *y*(*c*_*S*_(0)*− c*_*S*_(*t*)) = *m*(0)*− m*(*t*). Let us define the fibril mass concentration as *M* (*t*) = *m*(0)*− m*(*t*), i.e. the concentration of monomeric protein subunits aggregated into fibrils. We can then express the aggregation reaction yield as *M* (*∞*) = *m*(0) *− m*(*∞*).

Now, the concentration of SUVs is related to the lipid concentration *L*(*t*) as *c*_*S*_(*t*) = *L*(*t*)*/M*_*S*_, where *M*_*S*_ *≃* 6000 [28] is the average size of an SUV. So, the ratio of lipid to protein in the fibrils is (*L*(0) *− L*(*t*))*/*((*m*(0)*− m*(*t*)) = *M*_*S*_*/y*. The optimal stoichiometry of lipid to protein in lipidic fibrils, *x*, is estimated at 15, as discussed in Sec. . But the observed stoichiometry is not fixed judging by the slight negative curvature in the plot of yield vs *L*(0)*/m*(0) (Fig. 3). This demonstrates that the stoichiometry is slightly lowered when the initial ratio of lipid to monomeric protein *L*(0)*/m*(0) is below 15. This can be rationalized as the resultant improved yield resulting in lower free energy overall despite slightly less favourable free energy change per unit of fibril formed. Anyway, because all experiments analysed were performed at *L*(0)*/m*(0) well below 15, we estimate the stoichiometry as c. 15% lower than optimal, giving (*L*(0)*− L*(*t*))*/*((*m*(0)*− m*(*t*)) = 0.85*x* = *M*_*S*_*/y*. This gives *y ≃* 470, a value we use throughout all subsequent fitting.

The rate of nucleation appears to have no lipid dependence judging by Fig. 2, so we model nucleation processes in the normal way, writing for the total fibril end concentration *P* (*t*):

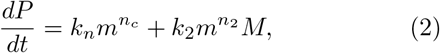

where the rate constants and reaction orders of primary and secondary nucleation are *k*_*n*_ and *n*_*c*_, and *k*_2_ and *n*_2_, respectively.

Since we know elongation involves lipid, and this is essentially entirely in the form of protein-coated SUVs, we must model both protein-addition and SUV-binding steps. Writing *P*_*i*_ as the concentration of fibril ends with SUVs bound *i* subunits away from the end, we now take a mean field approach to modelling. In this model, a monomer *cannot* bind to fibril ends with SUVs bound *y* subunits away from the end, or to the ends of freshly nucleated fibrils *P \**, and SUVs can *only* bind to fibril ends with SUVs bound *y* subunits away from the end, or to the ends of freshly nucleated fibrils. We also assume that monomers bind with the same rate to all fibril ends with SUVs bound closer than *y* subunits away. Thus,

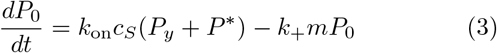

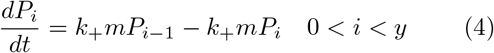

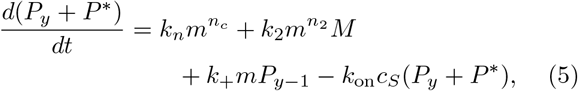

where *k*_+_ is the rate constant of elongation and *k*_on_ the rate constant of SUV addition. Summing these recovers Eq. (2) as required. Now, since nucleation rates are far slower than elongation rates, we can neglect nucleation and approximate a pre-equilibrium between these different types of fibril ends, leading to:

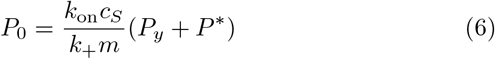

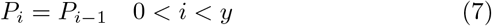

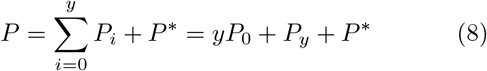

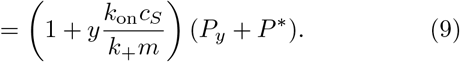

Now, the total rate of elongation is:

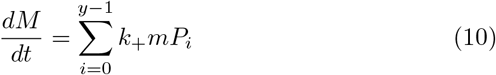

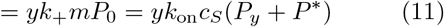

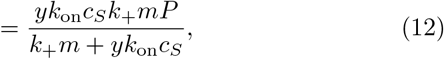

deriving Eq. (1b).

It should be fairly clear from the form of this equation that either monomer or lipid addition become rate-limiting depending on the relative values of *m* and *c*_*S*_. The cross-over between these regimes occurs at *k*_+_*m* = *yk*_on_*c*_*S*_ = *yk*_on_*L/M*_*S*_. We can rearrange this into the following constraint on rate constants:

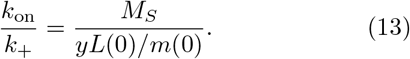

From Fig. 4 we see that the crossover occurs at *L*(0)*/m*(0) *≃* 4*/*3. Combined with *y ≃* 470 and *M*_*S*_ *≃* 6000, this gives *k*_on_*/k*_+_ *≃* 9.5. We use this constraint in all kinetic model fitting to prevent underdetermination.

### Kinetic analysis

Kinetic analysis is performed by numerically integrating the differential rate equations given above and using a least squares algorithm to fit the data. The data processing follows closely the procedure detailed in Meisl et al. [13].

### Secondary processes are only a minor contributor to formation of lipidic fibrils

To test whether secondary processes play a role in lipid-induced *α*-synuclein aggregation under our assay conditions, aggregation experiments were performed in which 10, 20, 30 and 40 _*μ*_M monomeric protein were incubated with DMPS vesicles at 5x and 8x lipid to protein molar ratios, and fibril formation was monitored by ThT fluorescence. Global fits to the model Eqs (1) were performed with *k*_2_ = 0 (Fig. 5**a-b**), or with *k*_2_ fitted along-side the other parameters (Fig. 5**c**-**d**). As expected, given the higher number of free fitting parameters, a non-zero secondary nucleation rate constant yielded improved fits. However, the difference in fit quality was not significant enough to positively confirm that secondary nucleation is present.

## Notes

### Competing Interest Statement

All authors were employees or consultants of WaveBreak Therapeutics when the relevant work was performed.

